# The cognitive effects of a promised bonus do not depend on dopamine synthesis capacity

**DOI:** 10.1101/2020.08.26.264168

**Authors:** Lieke Hofmans, Ruben van den Bosch, Jessica I. Määttä, Robbert-Jan Verkes, Esther Aarts, Roshan Cools

## Abstract

Reward motivation is known to enhance cognitive control. However, detrimental effects have also been observed, which have been attributed to overdosing of already high baseline dopamine levels by further dopamine increases elicited by reward cues. Aarts et al. (2014) indeed demonstrated, in 14 individuals, that reward effects depended on striatal dopamine synthesis capacity, measured with [^18^F]FMT-PET: promised reward improved Stroop control in low-dopamine individuals, while impairing it in high-dopamine individuals. Here, we aimed to assess this same effect in 44 new participants, who had previously undergone an [^18^F]DOPA-PET scan to quantify dopamine synthesis capacity. This sample performed the exact same rewarded Stroop paradigm as in the prior study. However, we did not find any correlation between reward effects on cognitive control and striatal dopamine synthesis capacity. The discrepancy between the current and our previous findings might reflect the use of different radiotracers for indexing dopamine synthesis capacity.

**STATEMENT OF RELEVANCE:** Reward motivation is generally thought to enhance cognitive control, but paradoxical negative effects of rewards on cognitive control have also been observed. A previous PET study demonstrated that reward effects on Stroop control depended on baseline striatal dopamine synthesis capacity, indexed by uptake of the radiotracer [^18^F]FMT. The sample size is this study was very small for a between-subject correlational design. Replicating the exact same Stroop paradigm within a larger sample is therefore crucial to robustly establish the mechanistic link between incentive motivation and cognitive control and advancing our understanding of who chokes under pressure and why, a topic of great societal relevance today. The present study did not reveal any correlation between reward effects on cognitive control and striatal dopamine synthesis capacity, indexed with [^18^F]FDOPA-PET. Future studies might consider putative differential sensitivity of the radiotracer [^18^F]FMT and [^18^F]FDOPA, while also addressing other indices of dopamine transmission.

## INTRODUCTION

Incentive (or reward) motivation is generally thought to enhance cognitive control [1, 2], However, paradoxical effects of rewards on cognitive control have also been observed, with some people exhibiting negative effects of reward anticipation on cognitive control [3–7], What mechanisms underlie these paradoxical effects of reward motivation on cognition? Motivational effects have long been associated with striatal dopamine signaling [8–10] and detrimental effects of rewards resonate with the notion of a potential overdosing of the dopamine system: Rewards, eliciting dopamine release [11], could have beneficial effects in individuals with low dopamine levels, but detrimental effects in individuals with already high dopamine levels [12], Prior work indeed suggested that effects of motivation on cognitive control depend on dopamine-related functioning, such as dopamine cell loss in Parkinson’s disease, midbrain and striatal BOLD activity, loss aversion and dopamine transporter genotype [3, 5, 6, 13, 14], Building on this work, our previous study [15] directly addressed this issue by assessing the effect of reward on cognitive control as a function of dopamine synthesis capacity, measured with 6-[^18^F]-fluoro-L-m-tyrosine ([^18^F]FMT) positron emission tomography (PET). Specifically, participants performed a Stroop task after being promised either a high or a low monetary reward upon successful completion of the task. These monetary incentives were demonstrated to impair Stroop interference control to a greater extent in participants with higher baseline dopamine synthesis capacity in the left caudate nucleus. It is of note that this effect was present only when participants were uninformed (i.e., un-cued) about the congruency of the upcoming Stroop target and not when Stroop targets were preceded by cues informing subjects about their congruency.

The finding that a negative correlation between reward effects and dopamine synthesis capacity was present specifically in the left caudate nucleus strengthened evidence from two other prior studies, implicating specifically the left caudate nucleus in the effects of the dopamine transporter gene *DAT1* during rewarded cognitive control [3, 16]. Moreover, this finding generally concurred with evidence from an fMRI study demonstrating enhanced connectivity between the ventral striatum and left caudate nucleus when cognitive demand for reward was high [17], The focus of the effect on the caudate nucleus also converged with functional MRI work from a third research group demonstrating a modulation by reward incentives of specifically the caudate nucleus during (oculomotor) control [18]. Finally, confidence in a negative correlation between individual differences in baseline dopamine levels and reward effects on cognitive control was further increased following a subsequent study in Parkinson’s disease patients, revealing greater beneficial effects of reward on cognitive control in patients with greater dopamine cell loss, measured with CIT-SPECT [13].

However, the sample size (n = 14) of the key PET study providing the direct evidence for baseline-dependency of reward effects on cognitive control in healthy volunteers was very small for a between-subject correlational design. Such a small effect size is associated not only with low positive predictive value [19], but also with high likelihood that effect sizes are biased and overestimated [20]. Therefore, we here aimed to test the effect found by Aarts *et al*. using a new, larger participant sample, who had already, as part of a previous study, undergone a PET scan with the radiotracer [^18^F]fluoro-3,4-dihydroxyphenyl-L-alanine ([^18^F]FDOPA). Specifically, we hypothesized that the effect of anticipated reward on Stroop interference control depends on individual differences in baseline dopamine synthesis capacity in the left caudate nucleus. We supplemented the analyses with voxel-wise correlations of the behavioral measures with dopamine synthesis capacity. Critically, striatal dopamine synthesis capacity in this new sample had already been indexed in the context of a previous study not reported here (www.trialregister.nl/trial/5959), using [^18^F]DOPA PET. This is unlike the original study, in which [^18^F]FMT PET was used to index striatal dopamine synthesis capacity.

The present attempt at conceptual replication was driven by our goal to increase our confidence in the role of dopamine synthesis capacity in motivated cognitive control and is of particular interest because a robust mechanistic account of the link between incentive motivation and cognitive control will advance our understanding of why and when who chokes under pressure [21], a topic of great societal relevance today. A preregistration of this study, data and code are available via https://osf.io/ky9s2/.

## MATERIALS AND METHODS

### Participants

Forty-five (out of a total of 94) right-handed and native Dutch-speaking volunteers who had participated in a previous [^18^F]DOPA PET study (protocol NL57538.091.16; trial register NTR6140, www.trialregister.nl/trial/5959) accepted the invitation to participate in the current study. All participants gave written informed consent according to the declaration of Helsinki and the experiment was conducted in compliance with and was approved by the local ethics committee (CMO Arnhem-Nijmegen, The Netherlands; Imaging Human Cognition, CMO 2014/288, version 2.2). One dataset was excluded due to an error rate above 33% (36%; mean (SD) = 18 (7) %). With the resulting 44 participants (aged: 19-45 years, mean (SD) = 24 (5.8); 22 women) we adhered to Simonsohn’s (2015) recommendation to obtain a sample size at least 2.5 times larger than the original sample size (N = 14). The new sample had 90% power [23] to detect a correlation of *r* = 0.55, which is considerably lower than the correlation of *r* = 0.75 reported in the original study (two-sided α = 0.0042, see Data analysis). The time between the PET scan and this behavioral study ranged between 0.3 and 1.8 years (mean (SD) = 1.0 (0.4)), which is substantially shorter than in the original study (range: 1.0-4.2 years, mean (SD) = 2.3 (1.1)). Background neuropsychological tests (listening span and behavioral inhibition / activation) had been assessed in the prior [^18^F]DOPA PET study.

### Behavioral paradigm

Participants completed the exact same paradigm as in Aarts *et al*.: a rewarded word-arrow Stroop paradigm, where they responded with a left or right button press to the words “left” or “right” in a left or right pointing arrow, using their right index finger or right middle finger, respectively (Figure 1A). The direction indicated by the word could either be congruent (same direction as the arrow) or incongruent (opposite direction). Each trial was preceded by a reward cue for a duration of 1-2 seconds, which indicated either a high (15 cents) or low (1 cent) reward that would be earned on that trial if the participant responded correctly and within the response window. After the reward cue, an information cue was shown on the screen for 1-2 seconds which was either informative, in which case it announced to the participant whether the trial would be congruent (green circle) or incongruent (red cross), or uninformative, in which case it showed a question mark. The information cues were added in the original study to assess potential anticipatory reward effects on proactive control, i.e. the ability to prepare for the upcoming congruent and incongruent Stroop targets (without being able to prepare a left or right motor response). Reward cues, information cues and congruency were equally divided across 240 trials, which lasted about 30 minutes.

**Figure 1.**
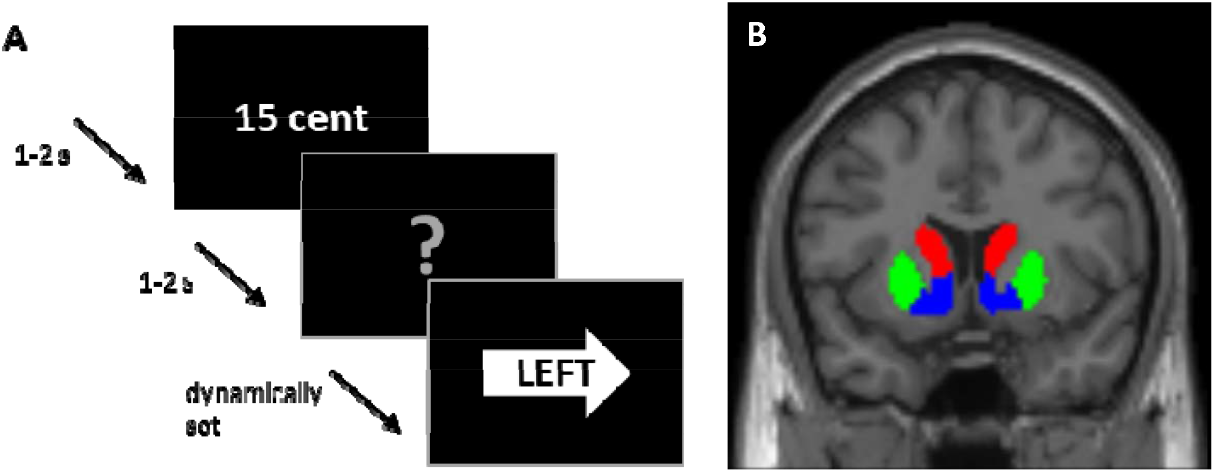
**A –** Schematic of an example word-arrow Stroop trial. Participants could either earn a high (15 cents) or low (1 cent) reward for a correct answer within the response window, which was cued at the start of the trial (1-2 seconds). After that, an information cue was presented for 1-2 seconds, indicating a congruent (green circle) or incongruent (red cross) trial, or giving no information about the upcoming congruency (grey question mark). Upon seeing the word-arrow Stroop target, participants had to respond to the word with a left or right button-press within the dynamically set response window. **(B)** Coronal view of our regions of interest including the bilateral caudate nucleus (red), putamen (green) and nucleus accumbens (blue).

As in the original study, before the actual task, participants completed 3 practice blocks. The first one to familiarize them with the information cues (12 trials), the second one to familiarize them with the reward cues (32 trials), and a third one - similar to the actual experiment - to set the initial response windows for the different trial types (48 trials). The initial response windows were set as the average response time per trial type (high or low reward; informed or uninformed; congruent or incongruent). During the actual experiment, the response windows were adapted after correct responses within the time window (−25 ms) or too late responses (+25 ms). After every block of 30 trials participants received feedback on their performance, showing their obtained reward on that block, the maximum reward that could have been obtained, their misses (too late), their errors (in time but wrong) and their reward for the total experiment so far, which remained on screen for 15 seconds. Due to the dynamic response windows, obtained reward was similar across participants and trial types (mean (SD) = €9.29 (0.94) for the entire experiment). The task was performed on a computer running on Windows 7 and a screen resolution of 1920×1080p, and the stimuli were shown using Presentation (version 20.2, 2018).

### PET acquisition

PET scans were carried out at the Department of Nuclear Medicine of the Radboud university medical center, using a Siemens PET/CT scanner and the radiotracer [^18^F]DOPA. Participants received 150 mg of carbidopa and 400 mg of entacapone 50 minutes before scanning, to minimize peripheral metabolism of [^18^F]DOPA and thereby increase central [^18^F]DOPA availability. The procedure started with a low dose CT scan (approximately 0.75 mCi) followed by a bolus injection of [^18^F]DOPA into an antecubital vein and an 89 minute dynamic PET scan (approximately 5 mCi). The data were divided into 24 frames (4×1, 3×2, 3×3, 14×5 min) and reconstructed with weighted attenuation correction and time-of-flight recovery, scatter corrected, and smoothed with a 3 mm full-width-at-half-maximum (FWHM) kernel.

### Structural MRI

A high-resolution anatomical scan, T1-weighted MP-RAGE sequence (repetition time = 2300 ms, echo time = 3.03 ms, 192 sagittal slices, field of view = 256 mm, voxel size 1 mm isometric) was acquired using a Siemens 3T MR scanner with a 64-channel coil. These were used for coregistration and spatial normalization of the PET scans.

### PET analysis

PET data were preprocessed and analyzed using SPM12 (http://www.fil.ion.ucl.ac.uk/spm/). All frames were realigned to the middle (11th) frame for motion correction and coregistered to the T1 MRI scan. Dopamine synthesis capacity was computed as the [^18^F]DOPA influx constant per minute (K_i_) per voxel relative to the grey matter of the cerebellum, using Gjedde-Patlak graphical analysis [24], The individual cerebellum grey matter masks were obtained by segmenting the individuals’ anatomical MRI scan, using Freesurfer (https://surfer.nmr.mgh.harvard.edu/). The K_i_ values were calculated based on the PET frames from the 24th to 89th minute. We then extracted average K_i_ values from six regions of interest (ROIs) - left and right caudate nucleus, putamen and nucleus accumbens - defined using masks based on an independent functional connectivity-analysis of the striatum (Piray et al., 2017; Figure 1B). For voxel-wise group analyses, the K¦maps were normalized to MNI space and smoothed using an 8 mm FWHM kernel.

### Data analysis

The main effect of interest was the correlation between the effect of reward on Stroop interference (in terms of response times) on uninformed trials and dopamine synthesis capacity in the left caudate nucleus, as was observed in the original study. For completeness, we also explored the other five ROIs. We analyzed response times (RTs) of all correct trials, including trials on which participants were “too late”, and error rates. Participants with error rates above 33% were excluded. We ran separate repeated measures analyses of variance (rmANOVA) for each region of interest and two dependent variables: Stroop interference on RT and on error rate (mean RT or error rate on incongruent trials minus mean RT or error rate on congruent trials). The within-subjects factors were REWARD (low, high) and INFORMATION (uninformed, informed), and [^18^F]DOPA K_i_ in the left or right caudate nucleus, putamen, or nucleus accumbens was a covariate of interest. The analyses were performed using the ezANOVA function from the ez package [26] in R (version 3.4.2). We corrected for multiple comparisons (6 ROIs, 2 dependent variables), resulting in a Bonferroni-corrected alpha value of 0.0042. Pearson’s correlations were calculated between the K_i_ values of the six ROIs and the effect of reward on Stroop interference in terms of RT on uninformed trials for comparison with the original study [15]. We supplemented the analyses with voxel-wise correlations between the reward effect on Stroop interference and dopamine synthesis capacity within the voxels comprising the entire striatum (the sum of the 6 regions of interest, specified above). Statistical significance was defined as family-wise error corrected *p* < 0.05 at peak coordinate, after small volume correction for all voxels within the striatal region of interest.

Although striatal [^18^F]DOPA uptake shows high test-retest reliability within a time frame of 2 years [27], we performed additional regression analyses, separately for each of the six ROIs, to assess whether any effects of the interaction between REWARD and dopamine synthesis capacity on Stroop interference depended on time between the PET scan and the behavioral testing day, while also including age and gender in the model, using the Im function from the stats package in R.

We could not directly compare baseline dopamine synthesis capacity between the original and the current study, because the PET tracer differed between the two studies. However, to appreciate possible differences between the main findings of the current study and that of the original study, it is important to analyze comparability of the sample (Table 1). We therefore compared the two samples in terms of age, neuropsychological assessment (listening span and behavioral inhibition / activation) and overall performance in terms of error rates and RT, using Welch’s t-tests. We then compared reward effects on Stroop interference in terms of RT between the two studies, with the hypothesis that reward would decrease interference in individuals with lower baseline dopamine synthesis capacity and increase interference in individuals with higher baseline dopamine synthesis capacity [12, 15]. We assessed differences in mean using a Welch’s t-test and differences in variances using a Levene’s test. Moreover, given the well-established link between dopamine and response vigor [9, 10, 28], we assessed the effect of baseline dopamine synthesis capacity on response times, both the main effect and in interaction with reward, in the current and the original study using an rmANOVA (Supplementary Table S4, S5).

**Table 1.** Demographic, background and task characteristics of participants included in the behavioral analyses.

|  | Characteristic | Aarts et al. (2014) | Current study | Welch's T | p |
| --- | --- | --- | --- | --- | --- |
| **Demographics** | Included participants | 14 | 44 |  |  |
|  | Sex (women) | 57% | 50% |  |  |
|  |  | **Mean (SD)** | **Mean (SD)** |  |  |
|  | Age (years) | 28 (2.7) | 24 (5.9) | -3.4 | 0.001 |
|  | Time between PET and behavioral testing (years) | 2.3 (1.1) | 1.0 (0.4) | -4.4 | 0.0006 |
| **Neuropsychological assessment** | Listening span |  |  |  |  |
|  | Total span | 3.8 (0.9)<sup>1</sup> | 4.3 (1.5) | 1.6 | 0.109 |
|  | Total words correct | 54.3 (7.7)<sup>1</sup> | 59.1 (16.0) | 1.5 | 0.143 |
|  | BIS/BAS |  |  |  |  |
|  | BIS | 19.5 (3.3) | 17.6 (4.3) | -1.8 | 0.087 |
|  | BAS (total score) | 37.4 (9.9) | 39.4 (4.2) | 0.76 | 0.457 |
| **Stroop task performance** | Overall error rate (%) | 15 (6) | 17 (7) | 1.0 | 0.319 |
|  | Overall response time (ms) | 398 (33) | 347 (57) | -4.1 | 0.0002 |
|  | Reward effect on Stroop interference on uninformed trials<sup>2</sup> | 0.1 (36.0) | -3.2 (32.3) | -0.3 | 0.767 |

Neuropsychological assessment included the listening span task [39] and the Behavioral Inhibition Scale/Behavioral Activation Scale (BIS/BAS; [40]). <sup>1</sup> 1 missing value; <sup>2</sup> average RT incongruent trials minus average RT congruent trials.

To allow for quantification of evidence for or against our hypotheses, we additionally report Bayesian individual effects analyses performed in JASP (version 0.10.2.0), with default JASP Cauchy priors. The BF_inclusion_ reflects how strongly the data support inclusion of a factor. We performed a sequential Bayesian correlation to illustrate evidence accumulation against the previously found correlation between the effect of reward on Stroop interference and dopamine synthesis capacity in the left caudate nucleus after observing the new data. Data from both studies were included; dopamine synthesis capacity values were separately standardized (z-scored) for both [^18^F]DOPA and [^18^F]FMT K_i_ values.

All continuous independent variables ([^18^F]DOPA Rvalues, time between the PET and behavioral session, overall response times and age) were mean centered.

## RESULTS

Participants performed more poorly on incongruent than congruent trials (RT: *F*_(1,43)_ = 185.3, *p* = 3.438e^-17^, BF_INC_ = 3.217e^+14^; error rate: *F*_(1,43)_ = 137.7, *p* = 5.394e^-15^, BF_INC_ = 6.434e^+13^), on uninformed than informed trials (RT: *F*_(1.43)_ = 35.0, *p* = 4.829e^-7^, BF_INC_ = 3.217e^+14^; error rate: *F*_(1,43)_ = 34.1, *p* = 6.224e^-7^, BF_INC_ = 4.949e^+13^) and low reward than high reward trials (RT: *F*_(1,43)_ = 14.2, *p* = 4.974e^-4^, BF_INC_ = 3.217e^+14^; error rate: *F*_(1,43)_ = 0.2, *p* = 0.662, BF_INC_ = 4.949e^+13^). These results validate the task manipulation, and they are similar to findings in the original study.

Crucially, and in contrast with the original study, there was no interaction effect between REWARD, INFORMATION and dopamine synthesis capacity in any of the six ROIs on Stroop interference in terms of response times or error rates (Table 2). For completeness, we also report the effect of reward on Stroop interference independent of the factor INFORMATION (Table 2). Pearson’s correlations between baseline dopamine synthesis capacity and the effect of reward on Stroop interference on uninformed trials only revealed no significant associations (all *r* < |0.22|, *p* > 0.158, BF < 0.575; Figure 2). Importantly, the 95% confidence interval for the correlation between dopamine synthesis capacity in the left caudate nucleus and the effect of reward on Stroop interference in the present study (*r* = −0.06, *p* = 0.700, 95% CI [-0.35, 0.24]) did not overlap with that of the originally reported effect of *r* = 0.75. Voxel-wise analyses of the effect of reward on Stroop interference on uninformed trials confirmed the lack of significant correlations with any of the voxels within the striatum (Figure 3). Separate multiple regression analyses for each ROI further confirmed the lack of a significant interaction between REWARD, INFORMATION and dopamine synthesis capacity or between REWARD and dopamine synthesis capacity on Stroop interference in terms of RT or error rate (Table 3). Additionally, time between PET and behavioral testing, age and gender did not affect the interaction between REWARD, INFORMATION and dopamine synthesis capacity or the interaction between REWARD and dopamine synthesis capacity on Stroop interference in terms of RT or error rate (Table 3). To further illustrate evidence against a correlation between the effect of REWARD on Stroop interference and dopamine synthesis capacity in the left caudate nucleus on uninformed trials, we ran a sequential Bayesian correlation including the data from both the original study and the current study. This revealed a strong increase in evidence in favor of a correlation when including participants from the original study, followed by a strong decline in evidence when including participants from the current study, culminating in moderate evidence against a correlation (Figure 4).

**Table 2.** Interaction effects in terms of response times (RT) and error rates obtained from the rmANOVAs with dopamine synthesis capacity in each ROI as a single covariate. The dependent variable is Stroop performance (mean RT or error rate on incongruent trials minus mean RT or error rate on congruent trials). Effect in grey was the interaction observed in Aarts *et al*. to be significant.

|  | Reward x information x DAsynth |  |  | Reward x DAsynth |  |  |
| --- | --- | --- | --- | --- | --- | --- |
| | $F_{(1,42)}$ | $p$ | $BF_{INC}$ | $F_{(1,42)}$ | $p$ | $BF_{INC}$ |
| <u>RT</u> |  |  |  |  |  |  |
| **Left caudate nucleus** | **1.9** | **0.177** | **0.003** | 0.5 | 0.473 | 0.044 |
| Right caudate nucleus | 0.6 | 0.456 | 0.003 | 1.4 | 0.244 | 0.041 |
| Left putamen | 2.4 | 0.126 | 0.003 | 0.2 | 0.628 | 0.029 |
| Right putamen | 3.4 | 0.072 | 0.004 | 0.3 | 0.612 | 0.029 |
| Left nucleus accumbens | 1.5 | 0.234 | 0.003 | 0.4 | 0.546 | 0.030 |
| Right nucleus accumbens | 0.8 | 0.365 | 0.002 | 0.0 | 0.964 | 0.034 |
| <u>Error rate</u> |  |  |  |  |  |  |
| Left caudate nucleus | 0.5 | 0.492 | 0.003 | 0.4 | 0.526 | 0.041 |
| Right caudate nucleus | 0.3 | 0.600 | 0.017 | 0.2 | 0.623 | 0.065 |
| Left putamen | 0.2 | 0.658 | 0.018 | 0.0 | 0.946 | 0.068 |
| Right putamen | 0.7 | 0.417 | 0.019 | 0.2 | 0.666 | 0.076 |
| Left nucleus accumbens | 0.0 | 0.951 | 0.014 | 0.3 | 0.618 | 0.059 |
| Right nucleus accumbens | 0.3 | 0.618 | 0.015 | 0.0 | 0.961 | 0.065 |

$p$ -values below a Bonferroni-corrected alpha-value of 0.0042 were considered significant.
Note that Aarts *et al.* analyzed the interaction between congruency, reward, information and dopamine synthesis capacity on response times and error rates. Here, we show the equivalent interaction between reward, information and dopamine synthesis capacity on Stroop interference (i.e. the difference between incongruent and congruent trials).

**Table 3.** Interaction effects obtained from multiple linear regression analyses assessing the effect of age, gender and time between the PET scan and behavioral testing (PET_behavioral) on motivational effects on Stroop interference (incongruent trials minus congruent trials) in terms of response times (RT) and error rates. Separate analysis for each ROI.

|  | Reward x<br>information x<br>DAsynth |  | Reward x<br>DAsynth |  | Reward x<br>information x<br>DAsynth x age |  | Reward x<br>DAsynth x age |  | Reward x<br>information x<br>DAsynth x<br>gender |  | Reward x<br>DAsynth x<br>gender |  | Reward x<br>information x<br>DAsynth x<br>PET_behavioral |  | Reward x<br>DAsynth x<br>PET_behavioral |  |
| --- | --- | --- | --- | --- | --- | --- | --- | --- | --- | --- | --- | --- | --- | --- | --- | --- |
| | $\beta$ | $p$ | $\beta$ | $p$ | $\beta$ | $p$ | $\beta$ | $p$ | $\beta$ | $p$ | $\beta$ | $p$ | $\beta$ | $p$ | $\beta$ | $p$ |
| <u>RT</u> |  |  |  |  |  |  |  |  |  |  |  |  |  |  |  |  |
| Left caudate nucleus | -3.4e<sup>3</sup> | 0.348 | -2.6e<sup>3</sup> | 0.481 | -3.7e<sup>2</sup> | 0.118 | -1.2e<sup>2</sup> | 0.629 | 3.3e<sup>3</sup> | 0.177 | 1.8e<sup>3</sup> | 0.449 | 5.5 | 0.592 | 4.7 | 0.644 |
| Right caudate nucleus | -3.6e<sup>3</sup> | 0.246 | -2.2e<sup>3</sup> | 0.472 | -2.1e<sup>2</sup> | 0.283 | -7.9e1 | 0.689 | 3.2e<sup>3</sup> | 0.134 | 1.4e<sup>3</sup> | 0.515 | 1.4 | 0.880 | -9.3e<sup>-2</sup> | 0.992 |
| Left putamen | 6.5e<sup>2</sup> | 0.845 | 1.5e<sup>3</sup> | 0.644 | 1.9e<sup>1</sup> | 0.945 | 6.3e<sup>1</sup> | 0.816 | 4.6e<sup>2</sup> | 0.837 | -4.4e<sup>2</sup> | 0.844 | 4.9 | 0.599 | 7.1 | 0.445 |
| Right putamen | 9.8e<sup>2</sup> | 0.756 | 1.6e<sup>3</sup> | 0.618 | 1.3e<sup>1</sup> | 0.959 | 7.4e<sup>1</sup> | 0.768 | 2.5e<sup>2</sup> | 0.900 | -5.2e<sup>2</sup> | 0.796 | 4.0 | 0.601 | 5.8 | 0.442 |
| Left nucleus accumbens | -2.0e<sup>3</sup> | 0.634 | 8.6e<sup>2</sup> | 0.833 | -5.6e<sup>2</sup> | 0.204 | -2.3e<sup>1</sup> | 0.959 | 1.5e<sup>3</sup> | 0.588 | -6.5e<sup>1</sup> | 0.981 | 1.3e<sup>1</sup> | 0.181 | 1.2e<sup>1</sup> | 0.230 |
| Right nucleus accumbens | -1.8e<sup>3</sup> | 0.614 | 8.1e<sup>2</sup> | 0.822 | -5.2e<sup>2</sup> | 0.082 | -1.4e<sup>2</sup> | 0.643 | 1.4e<sup>3</sup> | 0.551 | -2.5e<sup>2</sup> | 0.915 | 1.4e<sup>1</sup> | 0.123 | 1.4e<sup>1</sup> | 0.140 |
| <u>Error rate</u> |  |  |  |  |  |  |  |  |  |  |  |  |  |  |  |  |
| Left caudate nucleus | 6.6 | 0.697 | 1.5e<sup>1</sup> | 0.377 | -5.7e<sup>-2</sup> | 0.959 | -3.2e<sup>-1</sup> | 0.776 | -3.4 | 0.767 | -1.1e<sup>1</sup> | 0.322 | 2.1e<sup>-2</sup> | 0.661 | 8.8e<sup>-3</sup> | 0.855 |
| Right caudate nucleus | 1.1e<sup>1</sup> | 0.470 | 1.2e<sup>1</sup> | 0.430 | 5.4e<sup>-1</sup> | 0.570 | -7.8<sup>-1</sup> | 0.410 | -7.6 | 0.460 | -8.0 | 0.430 | 1.1e<sup>-2</sup> | 0.810 | 2.5e<sup>-2</sup> | 0.580 |
| Left putamen | 6.8e<sup>-2</sup> | 0.996 | 2.5e<sup>1</sup> | 0.100 | -8.2e<sup>-1</sup> | 0.513 | 2.2e<sup>-1</sup> | 0.859 | -1.9 | 0.854 | -1.7e<sup>1</sup> | 0.101 | 4.5e<sup>-4</sup> | 0.992 | -3.6e<sup>-3</sup> | 0.934 |
| Right putamen | 3.4 | 0.814 | 1.9e<sup>1</sup> | 0.195 | -5.9e<sup>-1</sup> | 0.608 | 3.6e<sup>-2</sup> | 0.975 | -4.8 | 0.601 | -1.3e<sup>1</sup> | 0.153 | -7.1e<sup>-3</sup> | 0.839 | -6.2e<sup>-3</sup> | 0.860 |
| Left nucleus accumbens | -1.1 | 0.955 | 1.9e<sup>1</sup> | 0.316 | -1.6 | 0.431 | 4.3e<sup>-1</sup> | 0.833 | -1.7 | 0.896 | -1.1e<sup>1</sup> | 0.379 | 1.2e<sup>-3</sup> | 0.979 | -3.8e<sup>-2</sup> | 0.408 |
| Right nucleus accumbens | -4.0 | 0.811 | 1.4e<sup>1</sup> | 0.401 | -9.8e<sup>-1</sup> | 0.478 | 3.9e<sup>-1</sup> | 0.778 | 2.3 | 0.837 | -8.6 | 0.442 | -9.8e<sup>-3</sup> | 0.820 | -2.2<sup>-2</sup> | 0.615 |

Model: stroop\_effect ~ DAsynth x reward x information x age + DAsynth x reward x information x gender + DAsynth x reward x information x PET\_behavioral

**Figure 2.**
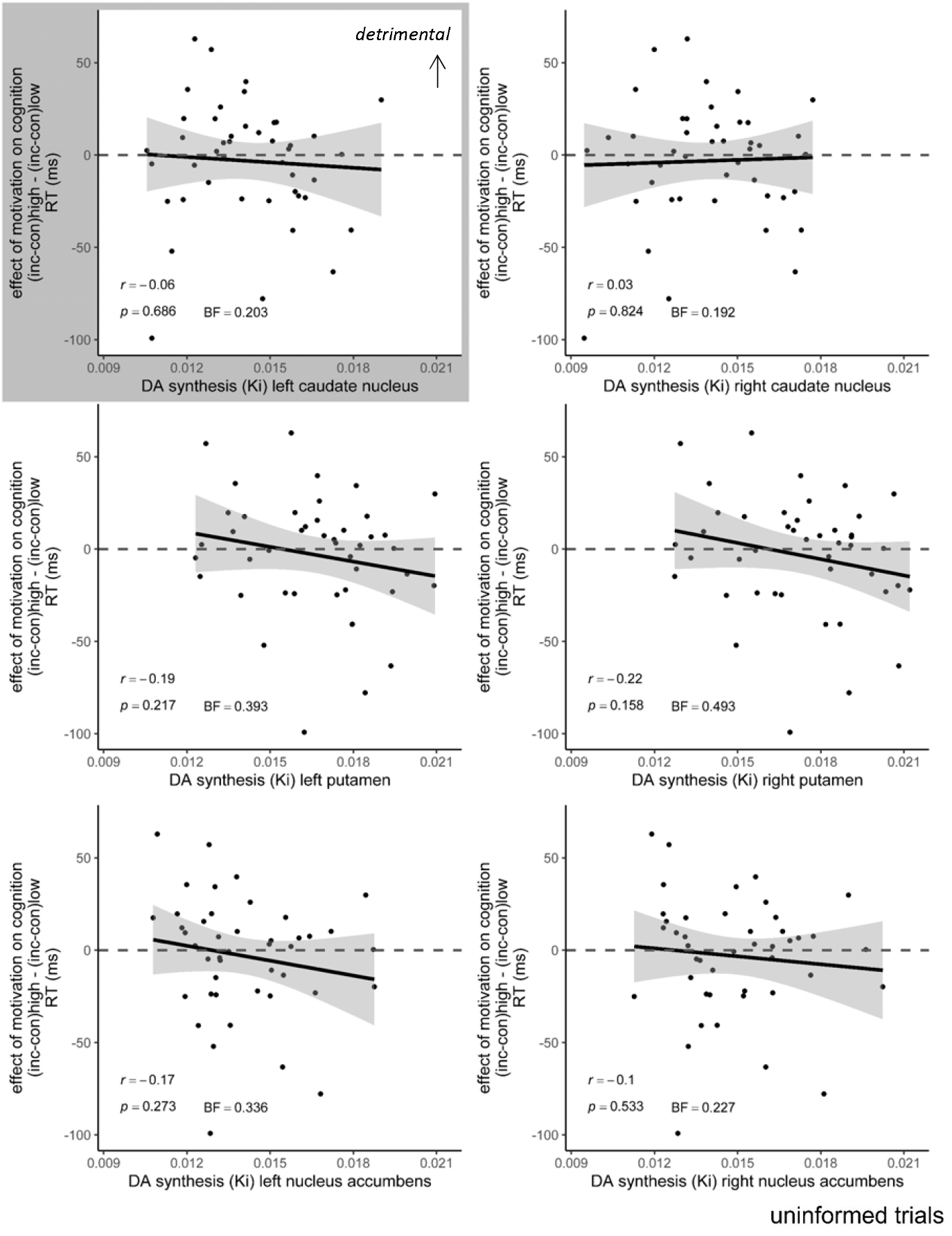
The effect of reward on Stroop interference (RT: incongruent – congruent) on uninformed trials plotted as a function of dopamine synthesis capacity in the six ROIs. Shaded area around the regression line represents 95% confidence interval. *N* = 44.; RT (ms) = response time in milliseconds; K_i_ = [^18^F]DOPA uptake, reflecting dopamine synthesis capacity. Effect in grey was the correlation observed in Aarts *et al*. to be significant.

**Figure 3.**
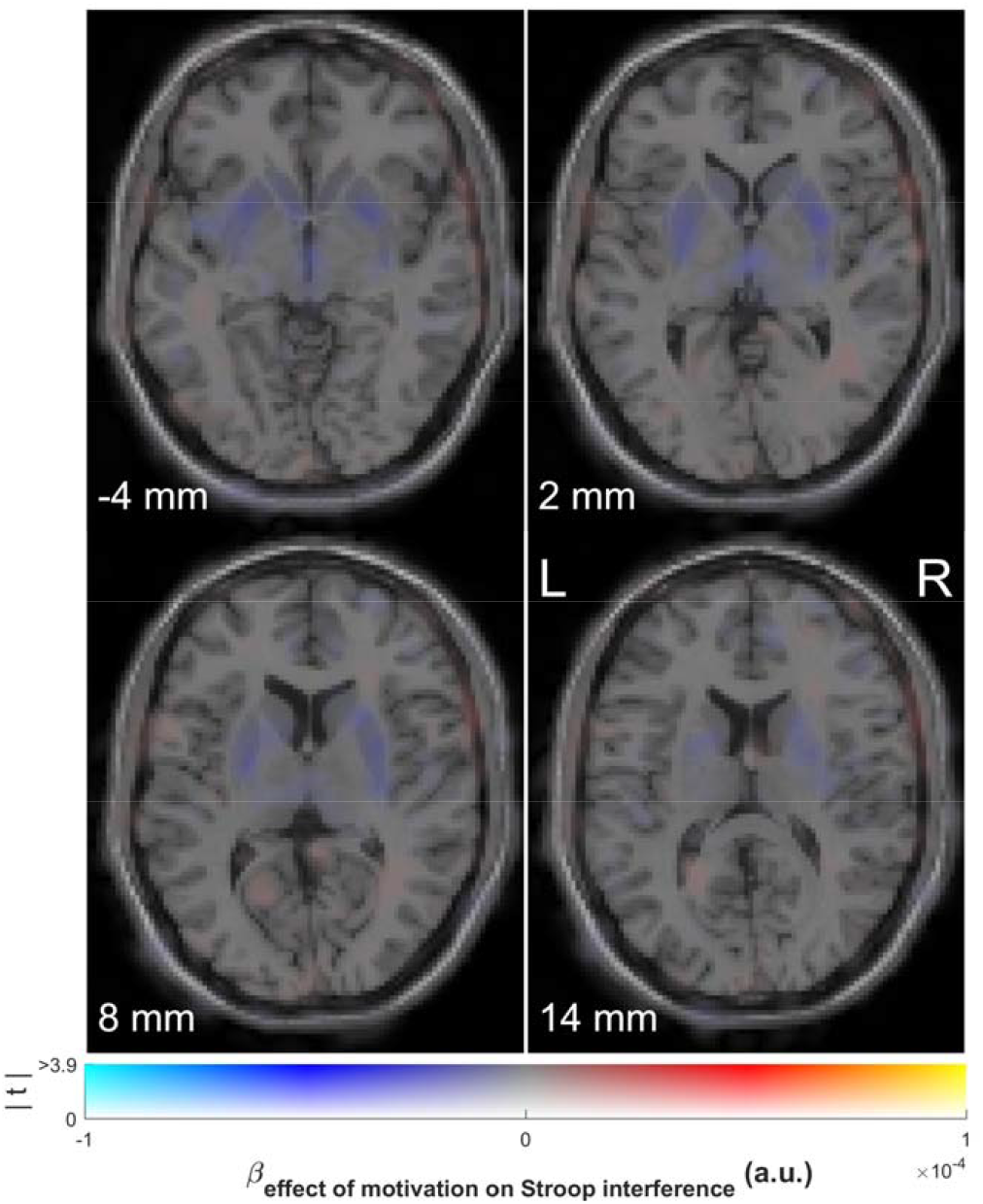
Association of baseline dopamine synthesis capacity with the effect of reward on Stroop interference on uninformed trials. Voxels showing a positive (red) or negative (blue) regression coefficient on the effect of a promised reward on Stroop interference in terms of response times on uninformed trials. Dual-coded and simultaneously displaying the contrast estimate (*x* axis) and *t* values (*y* axis). The hue indicates the size of the contrast estimate, and the opacity indicates the height of the *t* value. The *z* coordinates correspond to the standard MNI brain. No voxels survive *p* < 0.05 peak-level corrected (FWE) or *p* < 0.001 uncorrected. Plotted using a procedure introduced by Allen *et al*. [41] and implemented by Zandbelt [42].

**Figure 4.**
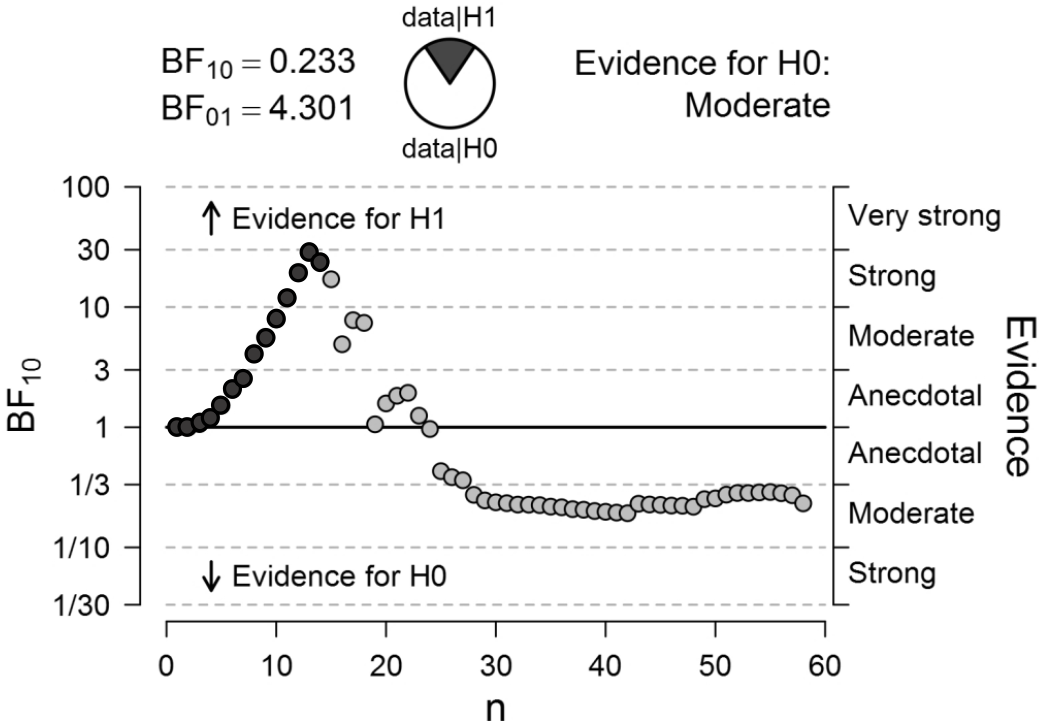
Sequential analysis showing progression of the Bayes Factor (BF) as new participants (n) enter the analysis. Values above 1 represent evidence for a correlation between dopamine synthesis capacity in the left caudate nucleus and a motivation effect on Stroop interference on uninformed trials, whereas values below 1 represent evidence against a correlation. Each dot represents the BF after inclusion of the next participant. Dark grey dots represent the 14 participants from *Aarts et al*. entered first; light grey dots represent the 44 participants from the current study. Order of including participants within each group was at random.

Average age differed significantly between the original and the current study (original study: mean = 28.1 years old; current study: mean = 24.3 years old; *t*_(47)_ = −3.4, *p* = 0.001; Table 1). To assess whether this could have caused the lack of effect of interest in the current study, we repeatedly discarded the youngest participant from our current dataset until age no longer differed between the studies, before rerunning the rmANOVAs. This resulted in a dataset including 26 participants (mean age = 26.9 years old; *t*_(37)_ = −0.9, *p* = 0.379). However, we did not observe a significant REWARD (by INFORMATION) by dopamine synthesis capacity interaction effect on Stroop interference (Supplementary Table S1).

Similarly, individual average RTs across trials differed significantly between the original and the current study (original study: mean = 397.5 ms; current study: mean = 346.9 ms; *t*_(40)_ = −4.1, *p* = 2.0e^-4^; Table 1). We therefore repeatedly discarded the fastest participant from our current dataset until the average RTs no longer differed, resulting in a dataset including 29 participants (mean RT = 371.8; *t*_(39)_ = −1.9, *p* = 0.064). However, we did not observe a significant REWARD (by INFORMATION) by dopamine synthesis capacity effect on Stroop interference (Supplementary Table S2). We additionally ran a multiple linear regression for each ROI including the terms REWARD, INFORMATION, dopamine synthesis capacity and individual average RT across all trials, including all interactions, which confirmed the lack of a significant effect of average RTs (Supplementary Table S3).

To establish that the discrepancy between the studies does not reflect differences in the dynamic range of the key variable of interest, we also compared the means and variances of the reward effects on Stroop interference on uninformed trials in terms of RT between the two studies. The two participant samples did not differ significantly from each other in terms of their means and variances (Figure 5), as revealed by a Welch’s t-test (original study: mean = 0.07 ms; current study: mean = −3.2 ms; *t*_(20)_ = −0.3, *p* = 0.767; Table 1) and Levene’s test (*F*_(1,56)_ = 0.2, *p* = 0.660), respectively.

**Figure 5.**
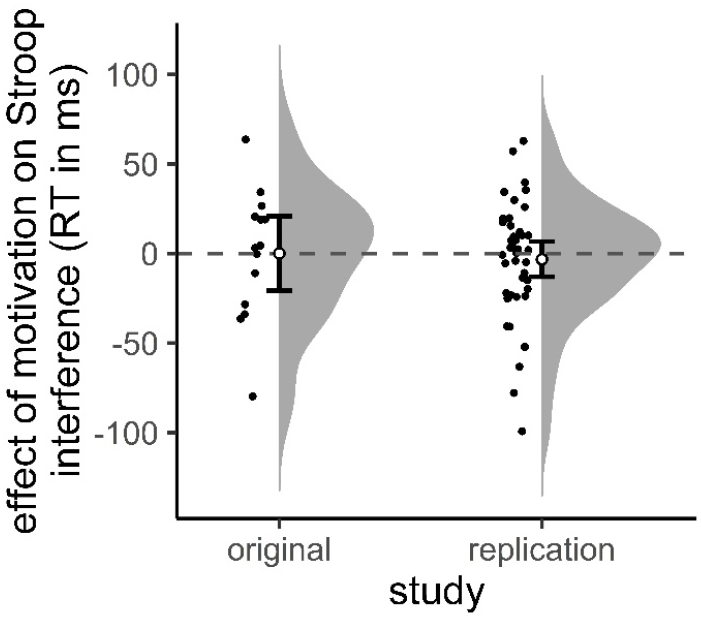
The effect of promised reward on Stroop interference (RT incongruent trials minus RT congruent trials) on uninformed trials in the original study (*Aarts et al*.) and the current replication attempt. Individual data points and probability distribution. Error bars represent 95% confidence interval around the mean. Plotted using R_rainclouds.R [43].

## DISCUSSION

The current study reveals no evidence for an interaction between monetary incentives and dopamine synthesis capacity, indexed with [^18^F]DOPA PET, on Stroop interference. Bayesian analyses in fact provide evidence in favor of a lack of a relationship between dopamine synthesis capacity and reward effect on Stroop interference. Our conclusion is therefore not consistent with the earlier findings by Aarts *et al*. [15].

It is possible that the discrepancy between the findings of the two studies reflects the use of [^18^F]DOPA in the present study, as opposed to [^18^F]FMT used in the original study. [^18^F]DOPA is a substrate for catechol-O-methyltransferase (COMT) in the periphery. Metabolites can cross the blood-brain-barrier and will distribute evenly throughout the brain, enhancing background noise relative to the use of [^18^F]FMT, which is not a substrate for COMT [29]. However, this is mainly a concern when one is interested in brain areas with low dopamine levels, as opposed to the dopamine-rich striatum. Moreover, entacapone was administered before PET scanning to inhibit peripheral COMT metabolism, further reducing the risk of a too low signal-to-noise ratio. [^18^F]DOPA and [^18^F]FMT also differ in their metabolic actions after decarboxylation by aromatic amino acid decarboxylase (AAAD), including higher affinity of [^18^F]DOPA metabolites compared with [^18^F]FMT metabolites for the vesicular monoamine transporter, leading to increased cell clearance of radiolabeled [^18^F]DOPA metabolites [30]. Indeed, differences in aging effects on dopamine synthesis capacity measured with [^18^F]DOPA and [^18^F]FMT have been observed, possibly owing to age-related changes in post-AADC metabolism [31]. However, this would mostly be a concern for extended scanning times, as [^18^F]DOPA behaves as an irreversibly bound tracer in the first 90 minutes after tracer injection, during which their uptake rates are tightly correlated [29, 30]. Another possibility is that the discrepancy between the original and the current study was introduced by group differences in sample characteristics. However, differences in overall response times and age did not explain the lack of significant effects in the current study. According to the dopamine overdose hypothesis [12], monetary incentives might enhance Stroop interference control in participants with very low average levels of baseline dopamine, whereas those incentives would impair control in participants with very high average levels. Sampling only participants with intermediate dopamine levels should lead to very small reward effects. However, a comparison of reward effects between the two studies demonstrated similar means and variances within the two samples. We therefore argue that the current result decreases our belief in the previously observed correlation between motivational effects on cognitive control and baseline dopamine synthesis capacity.

Notably, this conclusion would not imply that dopamine transmission is not important for the motivation of cognitive control, because brain dopamine levels are a function not only of dopamine synthesis capacity, but also of other factors, including transporter density, dopamine receptor availability, dopamine release and genetic make-up. Thus, the current study cannot refute hypothesized correlations between motivational effects on cognitive control and other measures of dopamine function. For example, the current design does not disconfirm previously demonstrated and replicated links between motivation, cognitive control and polymorphisms in the dopamine transporter gene [3, 14, 16], dopamine release [32] or dopamine-related disease status [13, 33, 34], Similarly, the current failure to replicate does not undermine other studies demonstrating a link between dopamine synthesis capacity and cognitive motivation indexed with other tasks, such as delay discounting [35], cognitive effort discounting [36, 37] or reward-based reversal learning [38]. Nevertheless, the presently observed lack of effect reduces our confidence in the link between dopamine synthesis capacity and the effect of a promised reward on Stroop interference and stresses the need for further studies.

## AUTHOR CONTRIBUTIONS

Conceptualization, R.C., L.H. and E.A.; Methodology E.A., R.C. and L.H.; Software, E.A. and L.H.; Formal analysis, L.H. and R.B.; Investigation, L.H.; Writing – Original Draft, L.H.; Writing – Review & Editing, L.H., R.C., E.A., R.-J.V., R.B., and J.I.M.; Visualization, L.H.; Supervision, R.C. and R.-J.V.; Project Administration, L.H. and J.I.M.; Funding Acquisition, R.C.

## ACKNOWLEDGMENTS AND DISCLOSURES

The work was funded by a Vici grant from the Netherlands Organization for Scientific Research (NWO) (Grant No. 453-14-015) awarded to RC. We thank Lieke van Lieshout and Felix Linsen for assistance during data collection.

## DATA AVAILABILITY

The data and analysis scripts used in this article will be made publicly available after manuscript acceptance at the following web address: https://data.donders.ru.nl/collections/published?4. Prior to accessing and downloading the shared data, users must create an account. It is possible to use an institutional account or a social ID from Google, Facebook, Twitter, LinkedIn or Microsoft. After authentication, users must accept the Data Use Agreement (DUA), after which they are automatically authorized to download the shared data. The DUA specifies whether there are any restrictions on how the data may be used. The Radboud University and the Donders Institute for Brain, Cognition and Behaviour will keep these shared data available for at least 10 years.

## Reference

1. Krawczyk DC, Gazzaley A, D’Esposito M. Reward modulation of prefrontal and visual association cortex during an incentive working memory task. Brain Res. 2007;1141:168–177.

2. Pessoa L, Engelmann JB. Embedding reward signals into perception and cognition. Front Neurosci. 2OIO;4.

3. Aarts E, Roelofs A, Franke B, Rijpkema M, Fernández G, Helmich RC, et al. Striatal dopamine mediates the interface between motivational and cognitive control in humans: Evidence from genetic imaging. Neuropsychopharmacology. 2010;35:1943–1951.

4. Aarts E, van Holstein M, Cools R. Striatal dopamine and the interface between motivation and cognition. Front Psychol. 2011;2:1–11.

5. Chib VS, De Martino B, Shimojo S, O’Doherty JP. Neural Mechanisms Underlying Paradoxical Performance for Monetary Incentives Are Driven by Loss Aversion. Neuron. 2012;74:582–594.

6. Mobbs D, Hassabis D, Seymour B, Marchant JL, Weiskopf N, Dolan RJ, et al. Choking on the Money. 2009;20:955–962.

7. Zedelius CM, Veling H, Aarts H. Boosting or choking - How conscious and unconscious reward processing modulate the active maintenance of goal-relevant information. Conscious Cogn. 2011;20:355–362.

8. Mohebi A, Pettibone JR, Hamid AA, Wong JMT, Vinson LT, Patriarchi T, et al. Dissociable dopamine dynamics for learning and motivation. Nature. 2019;570:65–70.

9. Robbins TW, Everitt BJ. A role for mesencephalic dopamine in activation: Commentary on Berridge (2006). Psychopharmacology (Berl). 2007;191:433–437.

10. Salamone JD, Yohn SE, Lo L. Activational and effort-related aspects of motivation: neural mechanisms and implications for psychopathology. Brain a J Neurol. 2016;139:1325–1347.

11. Schultz W. Dopamine neurons and their role in reward mechanisms. Curr Opin Neurobiol. 1997;7:191–197.

12. Cools R, D’Esposito M. Inverted-U-shaped dopamine actions on human working memory and cognitive control. Biol Psychiatry. 2011;69:e113–e125.

13. Aarts E, Helmich RC, Janssen MJR, Oyen WJG, Bloem BR, Cools R. Aberrant reward processing in Parkinson’s disease is associated with dopamine cell loss. Neuroimage. 2012;59:3339–3346.

14. van Holstein M, Aarts E, van der Schaaf ME, Geurts DEM, Verkes RJ, Franke B, et al. Human cognitive flexibility depends on dopamine D2 receptor signaling. Psychopharmacology (Berl). 2011;218:567–578.

15. Aarts E, Wallace DL, Dang LC, Jagust WJ, Cools R, D’Esposito M. Dopamine and the Cognitive Downside of a Promised Bonus. Psychol Sci. 2014;25:1003–1009.

16. Aarts E, van Holstein M, Hoogman M, Onnink M, Kan C, Franke B, et al. Reward modulation of cognitive function in adult attention-deficit/hyperactivity disorder. Behav Pharmacol. 2015;26:227–240.

17. Schmidt L, Lebreton M, Cléry-Melin ML, Daunizeau J, Pessiglione M. Neural mechanisms underlying motivation of mental versus physical effort. PLoS Biol. 2012;10.

18. Harsay HA, Cohen MX, Oosterhof NN, Forstmann BU, Mars RB, Ridderinkhof KR. Functional connectivity of the striatum links motivation to action control in humans. J Neurosci. 2011;31:10701–10711.

19. Heston TF, King JM. Predictive power of statistical significance. World J Methodol. 2017;7:112–116.

20. Button KS, Ioannidis JPA, Mokrysz C, Nosek BA, Flint J, Robinson ESJ, et al. Power failure: Why small sample size undermines the reliability of neuroscience. Nat Rev Neurosci. 2013;14:365–376.

21. Silston B, Mobbs D. Dopey dopamine: High tonic results in ironic performance. Trends Cogn Sci. 2014;18:340–341.

22. Simonsohn U. Small Telescopes: Detectability and the Evaluation of Replication Results. Psychol Sci. 2015;26:559–569.

23. Faul F, Erdfelder E, Buchner A, Lang AG. Statistical power analyses using G*Power 3.1: Tests for correlation and regression analyses. Behav Res Methods. 2009;41:1149–1160.

24. Patlak CS, Blasberg RG, Fenstermacher JD. Graphical evaluation of blood-to-brain transfer constants from multiple-time uptake data. Generalizations. J Cereb Blood Flow Metab. 1983;5:584–590.

25. Piray P, Ouden HEM Den, Schaaf ME Van Der, Toni I, Cools R. Dopaminergic Modulation of the Functional Ventrodorsal Architecture of the Human Striatum. 2017:485–495.

26. Lawrence MA. ez: Easy Analysis and Visualization of Factorial Experiments. R package version 4.4-0. https://CRAN.R-project.org/package=ez. 2016.

27. Egerton A, Demjaha A, McGuire P, Mehta MA, Howes OD. The test-retest reliability of 18F-DOPA PET in assessing striatal and extrastriatal presynaptic dopaminergic function. Neuroimage. 2010;50:524–531.

28. Niv Y, Daw ND, Joel D, Dayan P. Tonic dopamine: Opportunity costs and the control of response vigor. Psychopharmacology (Berl). 2007;191:507–520.

29. Becker G, Bahri MA, Michel A, Hustadt F, Garraux G, Luxen A, et al. Comparative assessment of 6-[18F]fluoro-L-m-tyrosine and 6-[18F]fluoro-L-dopa to evaluate dopaminergic presynaptic integrity in a Parkinson’s disease rat model. J Neurochem. 2017;141:626–635.

30. Doudet DJ, Chan GL-Y, Jivan S, DeJesus OT, McGeer EG, English C, et al. Evaluation of Dopaminergic Presynaptic Integrity: 6-[18 P]Fluoro-L-Dopa Versus 6-[ 18 P]Fluoro-L-m-Tyrosine. J Cerehral Blood Flow Metaholism. 1999;19:278–287.

31. DeJesus OT, Endres CJ, Shelton SE, Jerome Nickles R, Holden JE. Noninvasive assessment of aromatic L-amino acid decarboxylase activity in aging rhesus monkey brain in vivo. Synapse. 2001;39:58–63.

32. Jonasson LS, Axelsson J, Riklund K, Braver TS, Ögren M, Bäckman L, et al. Dopamine release in nucleus accumbens during rewarded task switching measured by [11C]raclopride. Neuroimage. 2014;99:357–364.

33. Manohar SG, Chong TTJ, Apps MAJ, Batla A, Stamelou M, Jarman PR, et al. Reward Pays the Cost of Noise Reduction in Motor and Cognitive Control. Curr Biol. 2015;25:1707–1716.

34. Timmer MHM, Aarts E, Esselink RAJ, Cools R. Enhanced motivation of cognitive control in Parkinson’s disease. Eur J Neurosci. 2018:0–2.

35. Smith CT, Wallace DL, Dang LC, Aarts E, Jagust WJ, D’Esposito M, et al. Modulation of impulsivity and reward sensitivity in intertemporal choice by striatal and midbrain dopamine synthesis in healthy adults. J Neurophysiol. 2016;115:1146–1156.

36. Hofmans L, Papadopetraki D, Bosch R van den, Määttä JI, Froböse MI, Zandbelt BB, et al. Baseline dopamine predicts individual variation in methylphenidate’s effects on cognitive motivation. BioRxiv. 2019:859637.

37. Westbrook A, van den Bosch R, Määttä J, Hofmans L, Papadopetraki D, Cools R, et al. Dopamine Promotes Cognitive Effort by Biasing the Benefits Versus Costs of Cognitive Work. BioRxiv Neurosci. 2019. 2019. https://doi.org/10.1101/778134.

38. Cools R, Frank MJ, Gibbs SE, Miyakawa A, Jagust W, D’Esposito M. Striatal Dopamine Predicts Outcome-Specific Reversal Learning and Its Sensitivity to Dopaminergic Drug Administration. J Neurosci. 2009;29:1538–1543.

39. Daneman M, Carpenter PA. Individual differences in working memory during reading. J Verbal Learning Verbal Behav. 1980;19:450–466.

40. Carver C, White T. Behavioral inhibition, behavioral activation, and affective responses to impending reward and punishment: The BIS/BAS Scales. J Pers Soc Psychol. 1994;67:319–333.

41. Allen EA, Erhardt EB, Calhoun VD. Data visualization in the neurosciences: overcoming the curse of dimensionality. Neuron. 2012;74:603–608.

42. Zandbelt B. Slice display, figshare. 2017:https://doi.org/10.%0A6084/m9.figshare.4742866.

43. Allen M, Poggiali D, Whitaker K, Marshall T, Kievit R. RainCloudPlots tutorials and codebase (Version v1.1). Zenodo. 2018.

