## Supplemental information for "The cognitive effects of a promised bonus do not depend on dopamine synthesis capacity"

Lieke Hofmans, Kapittelweg 29 r.2.269, 6525EN Nijmegen, The Netherlands

| **Table S1**. Interaction effects in terms of response times (RT) and error rates obtained from the rmANOVAs with dopamine synthesis capacity in each ROI as a single covariate. The dependent variable is Stroop performance (mean RT or error rate on incongruent trials minus mean RT or error rate on congruent trials). The sample is matched to the original sample from Aarts *et al.* in terms of age, resulting in N = 26. | | | | | | |
| --- | --- | --- | --- | --- | --- | --- |
|  | **Reward x information x DAsynth** | | | **Reward x DAsynth** | | |
|  | *F*_(1,24)_ | *p* | BF_INC_ | *F*_(1,24)_ | *p* | BF_INC_ |
| RT |  |  |  |  |  |  |
| Left caudate nucleus | 0.0 | 0.991 | 0.012 | 0.9 | 0.345 | 0.081 |
| Right caudate nucleus | 0.1 | 0.738 | 0.008 | 1.5 | 0.240 | 0.078 |
| Left putamen | 0.2 | 0.655 | 0.020 | 0.1 | 0.749 | 0.084 |
| Right putamen | 0.1 | 0.710 | 0.017 | 0.1 | 0.707 | 0.076 |
| Left nucleus accumbens | 0.0 | 0.931 | 0.015 | 0.1 | 0.737 | 0.070 |
| Right nucleus accumbens | 0.1 | 0.797 | 0.033 | 0.0 | 0.963 | 0.093 |
| Error rate |  |  |  |  |  |  |
| Left caudate nucleus | 0.1 | 0.806 | 0.121 | 0.1 | 0.717 | 0.231 |
| Right caudate nucleus | 0.2 | 0.683 | 0.180 | 0.0 | 0.848 | 0.281 |
| Left putamen | 1.5 | 0.236 | 0.351 | 0.0 | 0.879 | 0.369 |
| Right putamen | 2.4 | 0.138 | 0.391 | 0.3 | 0.583 | 0.432 |
| Left nucleus accumbens | 0.7 | 0.397 | 0.138 | 0.0 | 0.933 | 0.238 |
| Right nucleus accumbens | 0.2 | 0.674 | 0.218 | 0.0 | 0.931 | 0.323 |
| Note: *p*-values below a Bonferroni-corrected alpha-value of 0.0042 were considered significant. | | | | | | |

| **Table S2**. Interaction effects in terms of response times (RT) and error rates obtained from the rmANOVAs with dopamine synthesis capacity in each ROI as a single covariate. The dependent variable is Stroop performance (mean RT or error rate on incongruent trials minus mean RT or error rate on congruent trials). The sample is matched to the original sample from Aarts *et al.* in terms of individual average RT across all trials, resulting in N = 29. | | | | | | |
| --- | --- | --- | --- | --- | --- | --- |
|  | **Reward x information x DAsynth** | | | **Reward x DAsynth** | | |
|  | *F*_(1,27)_ | *p* | BF_INC_ | *F*_(1,27)_ | *p* | BF_INC_ |
| RT |  |  |  |  |  |  |
| Left caudate nucleus | 0.5 | 0.465 | 0.094 | 1.2 | 0.282 | 0.099 |
| Right caudate nucleus | 0.1 | 0.780 | 0.009 | 2.5 | 0.125 | 0.098 |
| Left putamen | 2.2 | 0.153 | 0.005 | 0.0 | 0.938 | 0.037 |
| Right putamen | 2.2 | 0.147 | 0.005 | 0.0 | 0.932 | 0.037 |
| Left nucleus accumbens | 0.6 | 0.459 | 0.004 | 0.0 | 0.935 | 0.039 |
| Right nucleus accumbens | 0.2 | 0.636 | 0.004 | 0.1 | 0.730 | 0.048 |
| Error rate |  |  |  |  |  |  |
| Left caudate nucleus | 0.2 | 0.651 | 0.016 | 0.0 | 0.884 | 0.041 |
| Right caudate nucleus | 0.0 | 0.884 | 0.014 | 0.0 | 0.945 | 0.054 |
| Left putamen | 2.8 | 0.106 | 0.028 | 0.2 | 0.697 | 0.082 |
| Right putamen | 5.6 | 0.025^1^ | 0.041 | 0.0 | 0.936 | 0.072 |
| Left nucleus accumbens | 0.4 | 0.551 | 0.013 | 0.5 | 0.490 | 0.073 |
| Right nucleus accumbens | 0.4 | 0.517 | 0.014 | 0.0 | 0.954 | 0.085 |
| Note: *p*-values below a Bonferroni-corrected alpha-value of 0.0042 were considered significant.  ^1^*p*-value does not survive correction for multiple comparisons. However, for clarity we report the interaction effect of reward x DAsynth in the right Putamen on Stroop interference (error rate): informed trials: *r* = -0.21, *p* = 0.266; uninformed trials: *r* = 0.29, *p* = 0.131. | | | | | | |

| **Table S3**. Interaction effects obtained from multiple linear regression analyses assessing the effect of individual average RT across all trials on motivational effects on Stroop interference (incongruent trials minus congruent trials) in terms of response times (RT) and error rates. Separate analysis for each ROI. | | | | | | | | |
| --- | --- | --- | --- | --- | --- | --- | --- | --- |
|  | **Reward x information x DAsynth** | | **Reward x**  **DAsynth** | | **Reward x information x DAsynth x RT** | | **Reward x**  **DAsynth x RT** | |
|  | β | *p* | β | *p* | β | *p* | β | *p* |
| RT |  |  |  |  |  |  |  |  |
| Left caudate nucleus | 5.2e^3^ | 0.207 | -7.3e^3^ | 0.260 | -5.9e^1^ | 0.416 | 1.3e^2^ | 0.251 |
| Right caudate nucleus | 4.7e^3^ | 0.260 | -6.3e^3^ | 0.330 | -4.2e^1^ | 0.510 | 1.1e^2^ | 0.280 |
| Left putamen | 4.4e^3^ | 0.248 | -7.0e^3^ | 0.249 | 1.8e^1^ | 0.870 | 1.3e^1^ | 0.939 |
| Right putamen | 4.5e^3^ | 0.219 | -7.1e^3^ | 0.223 | -2.7e^1^ | 0.797 | 8.8e^1^ | 0.602 |
| Left nucleus accumbens | 3.8e^3^ | 0.371 | -6.5e^3^ | 0.334 | -2.1e^1^ | 0.819 | 7.2e^1^ | 0.819 |
| Right nucleus accumbens | 3.1e^3^ | 0.424 | -5.1e^3^ | 0.405 | -5.1e^1^ | 0.486 | 1.1e^2^ | 0.333 |
| Error rate |  |  |  |  |  |  |  |  |
| Left caudate nucleus | 3.0 | 0.878 | 2.5 | 0.934 | 1.1e^-1^ | 0.738 | 6.3e^-2^ | 0.906 |
| Right caudate nucleus | -3.8 | 0.846 | 1.2e^1^ | 0.699 | 2.4e^-1^ | 0.419 | -1.3e^-1^ | 0.780 |
| Left putamen | -8.2 | 0.648 | 1.4e^1^ | 0.622 | -2.2e^-1^ | 0.661 | 5.4e^-1^ | 0.501 |
| Right putamen | -1.2e^1^ | 0.505 | 2.2e^1^ | 0.416 | -1.8e^-1^ | 0.717 | 4.9e^-1^ | 0.535 |
| Left nucleus accumbens | -3.8e^-2^ | 0.998 | -2.1 | 0.947 | 1.0e^-1^ | 0.814 | 1.0e^-1^ | 0.881 |
| Right nucleus accumbens | 3.8 | 0.834 | -5.6 | 0.847 | 2.1e^-3^ | 0.995 | 2.5e^-1^ | 0.650 |
| Model: stroop_effect ~ DAsynth x reward x information x average RT | | | | | | | | |

Both in the current sample (Table S4) and the original sample (Table S5), baseline dopamine synthesis capacity was not associated with response times, neither in interaction with reward nor as main effect:

| **Table S4**. Effect of dopamine synthesis capacity and dopamine synthesis capacity x reward on response times in the current sample. Separate rmANOVA for each of the six regions of interest, including reward, congruency and information as within-subjects factors and dopamine synthesis capacity as covariate. Higher N = 44. | | | | |
| --- | --- | --- | --- | --- |
|  | **DAsynth** | | **Reward x DAsynth** | |
|  | *F*_(1,42)_ | *p* | *F*_(1,42)_ | *p* |
| Left caudate nucleus | 2.4 | 0.128 | 3.1 | 0.084 |
| Right caudate nucleus^1^ | 7.5 | 0.009 | 3.8 | 0.058 |
| Left putamen | 0.0 | 0.833 | 1.1 | 0.299 |
| Right putamen | 0.1 | 0.804 | 1.3 | 0.268 |
| Left nucleus accumbens | 0.1 | 0.771 | 1.3 | 0.261 |
| Right nucleus accumbens | 0.3 | 0.596 | 2.9 | 0.095 |
| ^1^Corresponds to a negative correlation between dopamine synthesis capacity and response times. Effects when 1 participant with low dopamine synthesis capacity in the right caudate nucleus and an average RT of 4 standard deviations above the group mean was excluded: DAsynth: *F*_(1,41)_ = 3.0, *p* = 0.092; reward x DAsynth: *F*_(1,41)_ = 0.4, *p* = 0.535. | | | | |

| **Table S5**. Effect of dopamine synthesis capacity and dopamine synthesis capacity x reward on response times in Aarts *et al*. Separate rmANOVA for each of the six regions of interest, including reward, congruency and information as within-subjects factors and dopamine synthesis capacity as covariate. N = 14. | | | | |
| --- | --- | --- | --- | --- |
|  | **DAsynth** | | **Reward x DAsynth** | |
|  | *F*_(1,12)_ | *p* | *F*_(1,12)_ | *p* |
| Left caudate nucleus | 0.1 | 0.738 | 0.0 | 0.930 |
| Right caudate nucleus | 0.0 | 0.855 | 0.0 | 0.871 |
| Left putamen | 0.0 | 0.966 | 0.0 | 0.965 |
| Right putamen | 0.0 | 0.907 | 0.1 | 0.817 |
| Left nucleus accumbens | 0.2 | 0.702 | 0.0 | 0.893 |
| Right nucleus accumbens | 0.1 | 0.780 | 0.3 | 0.567 |
